# Antiviral defense via nucleotide depletion in bacteria

**DOI:** 10.1101/2021.04.26.441389

**Authors:** Nitzan Tal, Adi Millman, Avigail Stokar-Avihail, Taya Fedorenko, Azita Leavitt, Sarah Melamed, Erez Yirmiya, Carmel Avraham, Gil Amitai, Rotem Sorek

## Abstract

DNA viruses and retroviruses need to consume large quantities of deoxynucleotides (dNTPs) when replicating within infected cells. The human antiviral factor SAMHD1 takes advantage of this vulnerability in the viral life cycle, and inhibits viral replication by degrading dNTPs into their constituent deoxynucleosides and inorganic phosphate. In this study we report that bacteria employ a similar strategy to defend against phage infection. We found a family of defensive dCTP deaminase proteins that, in response to phage infection, convert dCTP into deoxy-uracil nucleotides. A second family of phage resistance genes encode dGTPase enzymes, which degrade dGTP into phosphate-free deoxy-guanosine (dG) and are distant homologs of the human SAMHD1. Our results show that the defensive proteins completely eliminate the specific deoxynucleotide (either dCTP or dGTP) from the nucleotide pool during phage infection, thus starving the phage of an essential DNA building block and halting its replication. Both defensive genes are found in a diverse set of bacterial species and are specifically enriched in *Vibrio* genomes. Our study demonstrates that manipulation of the deoxynucleotide pool is a potent antiviral strategy shared by both prokaryotes and eukaryotes.

## Introduction

Bacteria encode multiple mechanisms of immunity to defend themselves against phage infection (Bernheim and Sorek, 2020; Hampton et al., 2020). Restriction-modification (RM) and CRISPR-Cas systems have long been recognized as major lines of defense against phage (Hampton et al., 2020). Abortive infection systems, which lead infected cells to commit suicide and thus abort phage propagation, have also been documented in bacteria since the early days of phage research (Lopatina et al., 2020).

In recent years, it has become clear that bacteria encode a plethora of additional immune systems that escaped early detection (Bernheim and Sorek, 2020; Hampton et al., 2020). These include defense systems that produce small molecules which block phage replication (Bernheim et al., 2021; Kronheim et al., 2018), systems that rely on secondary-messenger signaling molecules that activate immune effectors (Cohen et al., 2019; Lau et al., 2020; Lowey et al., 2020; Ofir et al., 2021), and retron systems that employ reverse-transcribed non-coding RNAs as part of their anti-phage activity (Bobonis et al., 2020; Millman et al., 2020). For the vast majority of newly discovered defense systems, the mechanism is still unknown, and it has been hypothesized that many bacterial defense systems still await discovery (Gao et al., 2020).

Bacterial defense systems are commonly localized in genomic ‘defense islands’, a characteristic that has facilitated the discovery of many new systems based on their tendency to co-localize with known defense systems such as RM and CRISPR-Cas (Doron et al., 2018; Gao et al., 2020). Another distinctive feature of bacterial defense systems is their rapid pace of loss and re-gain by horizontal gene transfer over short time scales, so that even in closely related strains that have otherwise similar genomes, the composition of defense systems can vary substantially (Bernheim and Sorek, 2020).

In this study, we describe two previously unknown families of bacterial defense proteins that, in response to phage infection, eliminate one of the deoxynucleotides (either dCTP or dGTP) from the nucleotide pool, thus depriving the phage from an essential DNA building block and halting its replication. One of these families, encoding dGTP-degrading enzymes, shows homology to the human SAMHD1 antiviral protein which similarly defends against HIV and other viruses by degrading dNTPs to deplete the nucleotide pool. Our study demonstrates convergence of antiviral immune mechanisms from bacteria to humans.

## Results

### A family of anti-phage cytidine deaminases

We initiated this study by focusing on a family of 977 homologous genes that had a predicted cytidine deaminase domain. This family of genes caught our attention due to their frequent localization next to known anti-phage defense systems in diverse bacterial genomes (Figure 1A-1B; Supplementary Table S1). Such preferential genomic localization near defense systems was previously shown to be a strong predictor for a role in anti-phage defense (Doron et al., 2018) and we therefore hypothesized that these cytidine deaminase genes similarly had an anti-phage function. To test this hypothesis, we cloned two genes from this family, one from *Escherichia coli* U09 and the other from *E. coli* AW1.7, into a lab strain of *E. coli* (MG1655) that does not naturally encode such genes. Infection assays with a panel of phages showed that both genes conferred substantial defense against a diverse set of phages (Figure 1C; Figure S1). Since it conferred wider defense against phages, we further functionally characterized the gene from *E. coli* AW1.7 (Figure 1C-1D; Figure S1).

**Figure 1.**
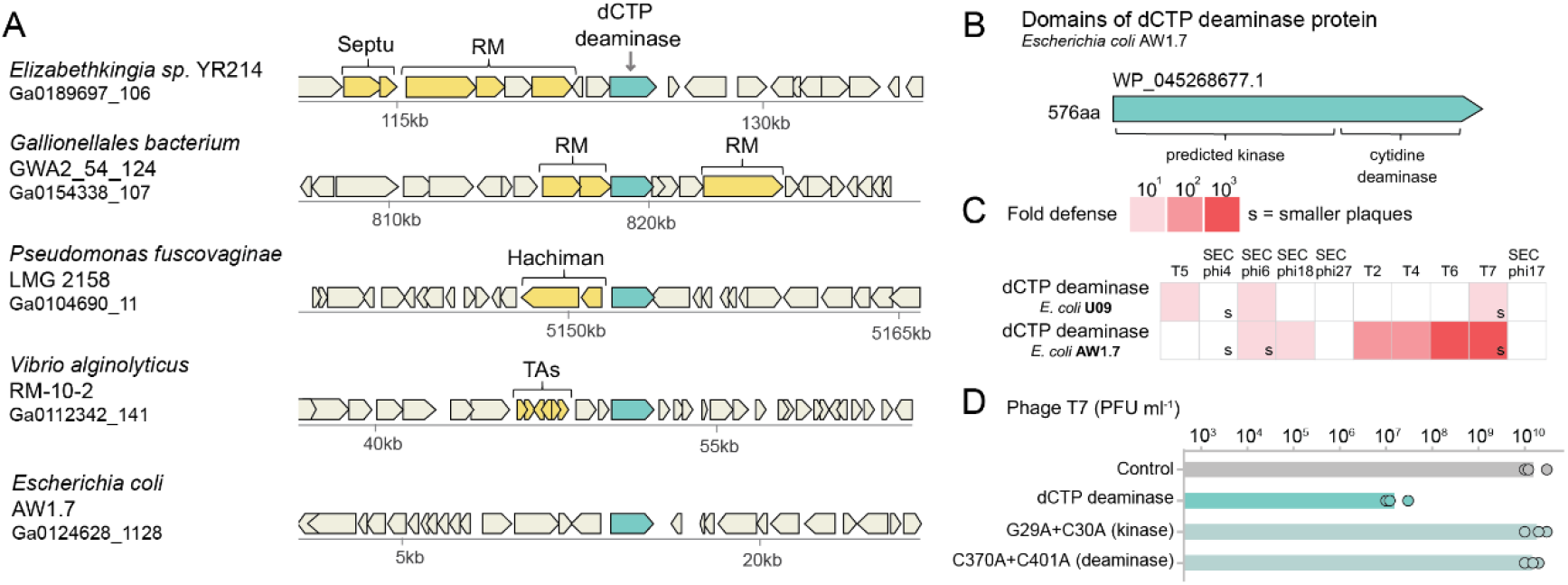
A family of cytidine deaminases provide defense against phages. (A) Representative instances of cytidine deaminase genes (in turquoise) and their genomic neighborhoods. Genes known to be involved in defense are shown in yellow. RM, restriction modification; TAs, toxin-antitoxin systems; Septu and Hachiman are recently described defense systems (Doron et al., 2018). The bacterial species and the accession of the relevant genomic scaffold in the Integrated Microbial Genomes (IMG) database (Chen et al., 2019) are indicated on the left. (B) Domain organization of the cytidine deaminase of *E. coli* AW1.7. Protein accession in NCBI is indicated above the gene. (C) Cytidine deaminases defend against phages. Cytidine deaminases from two *E. coli* strains were cloned, together with their promoter regions, and transformed into *E. coli* MG1655. Fold defense was measured using serial dilution plaque assays, comparing the efficiency of plating (EOP) of phages on the deaminase-containing strain to the EOP on a control strain that lacks the gene. Data represent an average of three replicates (see detailed data in Figure S1). The designation ‘s’ stands for a marked reduction in plaque size. (D) Effect of point mutations on the defensive activity of dCTP deaminase from *E. coli* AW1.7. Data represent plaque-forming units per ml (PFU/ml) of T7 phages infecting control cells, dCTP deaminase-expressing cells, and two strains mutated in the predicted kinase or deaminase domains. Shown is the average of three replicates, with individual data points overlaid.

Cytosine deaminase enzymatic activities, which convert cytosine into uracil bases, have been recorded in multiple immune proteins that protect human cells from viral infection (Harris and Dudley, 2015). Specifically, the human protein APOBEC3G protects against retroviruses by inducing numerous deoxycytidine-to-deoxyuridine substitutions in the complementary DNA of the viral genome, causing hyper-mutations that destroy the coding capacity of the virus (Harris and Dudley, 2015). Based on this, we initially hypothesized that the cytosine deaminase genes we found in bacteria may protect against phages by introducing mutations into the viral DNA or RNA.

To examine this hypothesis, we infected the deaminase-containing and control strains with phage T7 and extracted total DNA and RNA from the infected cells at several time points following initial infection. We then sequenced the extracted nucleic acids and aligned the sequences to the viral and host genomes, searching for C-to-T mismatches (or G-to-A in the complementary strand). We could not find a difference between the rates of C-to-T or G-to-A mismatches in phage DNA or RNA in *E. coli* MG1655 cells encoding the cytidine deaminase as compared to control cells lacking the gene (Figure S2). In addition, no hyper-editing of cytosines in the DNA or RNA molecules of the bacterial host was observed (Figure S2). We also could not find any positions in the phage or the host genomes and transcriptomes that were specifically converted from cytosine to uracil or thymine. We therefore concluded that the bacterial cytosine deaminase genes protect against phages via a mechanism that does not involve editing of the nucleic acid polymers.

### Deamination of dCTP depletes it from the nucleotide pool

We next examined the possibility that the bacterial defensive protein modifies single nucleotides rather than DNA or RNA polymers. To this end, we filtered extracts from cells infected by phage T7 and used liquid chromatography followed by mass spectrometry (LC-MS) to monitor the nucleotide content in these cell extracts (Figure 2A). Remarkably, deoxycytidine triphosphate (dCTP), which naturally accumulates in control cells infected by phage T7, was completely absent in infected cells expressing the deaminase gene (Figure 2B). The depletion of dCTP was associated with substantial elevation of deoxy-uridine monophosphate (dUMP), recorded as early as five minutes after initial infection, suggesting that the observed loss of dCTP was caused by its deamination into deoxy-uridine compounds (Figure 2C). Similar depletion was observed in deoxycytidine monophosphate (dCMP) and deoxycytidine diphosphate (dCDP), but not in the ribonucleotides CTP, CDP, and CMP, suggesting that deamination is limited to deoxycytidines (Figures 2D-2E; Figure S3). Although the deamination of dCTP is expected to generate dUTP molecules, it is known that *E. coli* expresses housekeeping dUTPase enzymes that rapidly convert dUTP to dUMP molecules (Vértessy and Tóth, 2009), likely explaining the observed accumulation of dUMP and not of dUTP (Figure 2C).

**Figure 2.**
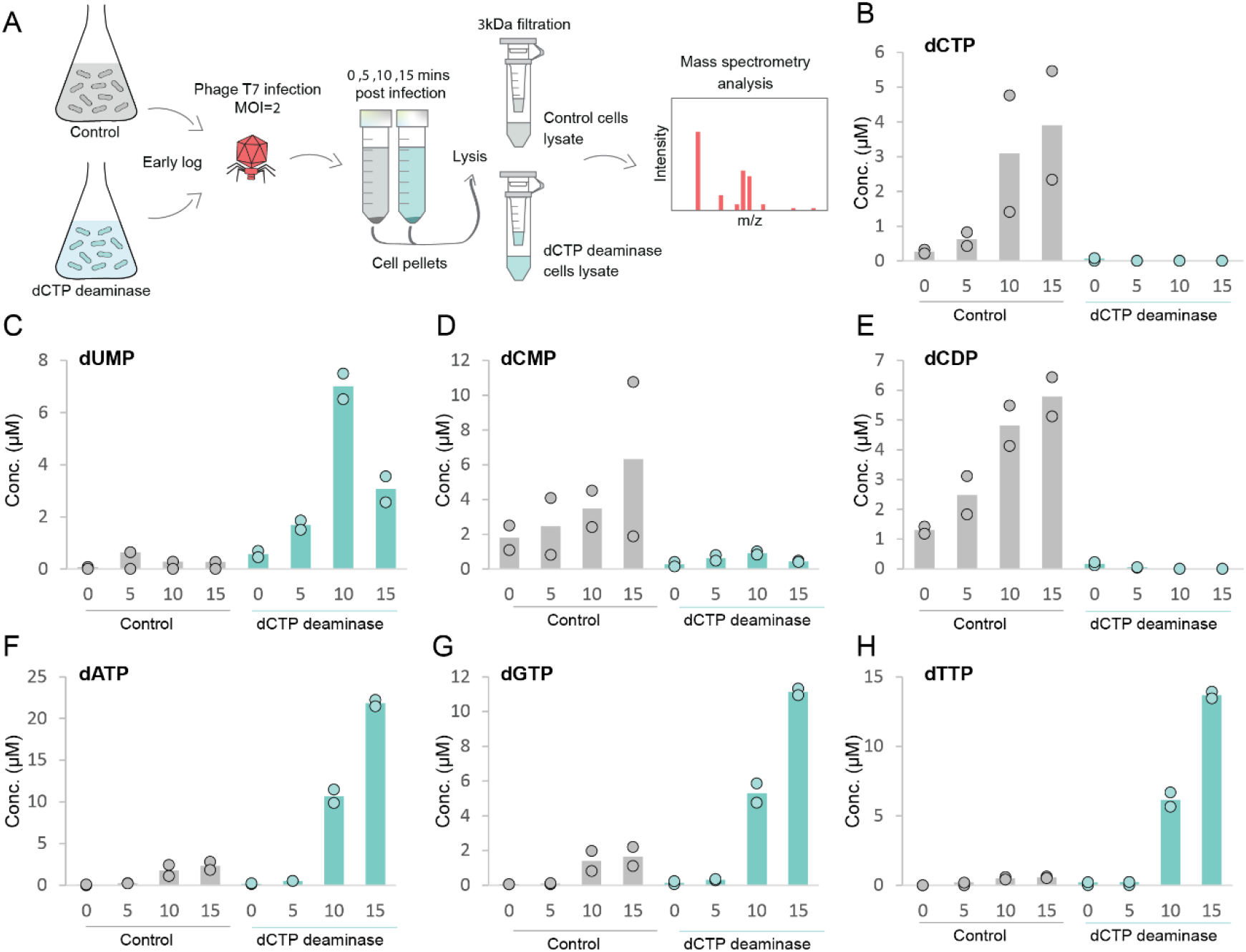
Depletion of deoxycytidine nucleotides during phage infection. (A) Schematic representation of the experiment. (B-H) Concentrations of deoxynucleotides in cell lysates extracted from T7-infected cells, as measured by LC-MS with synthesized standards. X-axis represents minutes post infection, with zero representing non-infected cells. Cells were infected by phage T7 at an MOI of 2 at 37°C. Each panel shows data acquired for dCTP deaminase-expressing cells or for control cells that contain an empty vector. Bar graphs represent the average of two biological replicates, with individual data points overlaid.

These results suggest that during infection by phage T7, the defensive deaminase protein converts deoxycytidine nucleotides into deoxyuracils, depleting the cell of the dCTP building blocks that are essential for phage DNA replication. In support of this hypothesis, we observed that the other three deoxy nucleotides, dATP, dGTP and dTTP, all accumulated in the cell at 10 and 15 minutes post infection (Figures 2F-2H). The concentration of these nucleotides also naturally increases in control cells infected by T7, since the phage produces them by breaking down the host DNA to serve as building blocks for its replicating genome (Lee and Richardson, 2011); however, in cells expressing the defensive deaminase, levels of dATP, dGTP and dTTP further increased more than five-fold as compared to control cells (Figures 2F-2H). It is likely that the absence of dCTP prevents DNA replication, resulting in the observed buildup of the nucleotide building blocks that would have otherwise been incorporated into the replicating polynucleotide chain of the phage. Indeed, dNTP accumulation was previously observed in T7-infected cells in which DNA synthesis was blocked (Myers et al., 1987).

The dCTP deaminase gene that we studied encodes a 576-amino-acid protein. Its C-terminus encompasses a predicted zinc-dependent cytosine deaminase domain, while the N-terminus is occupied by a predicted kinase domain (Figure 1B). Point mutations in the active site of the deaminase domain (C370A and C401A, predicted to disrupt zinc binding) abolished defense (Figure 1D). In addition, point mutations in the predicted nucleotide-binding motif of the kinase domain (G29A and C30A) also abolished defense (Figure 1D). Accordingly, dCTP was not depleted during phage infection in cells expressing either the deaminase or the kinase mutants (Figure S4). These results suggest that both the deaminase and the kinase domains are essential for the ability of the dCTP deaminase to defend via dCTP depletion.

### Defensive dGTPases deplete dGTP during phage infection

Our discovery of a new defensive mechanism that employs nucleotide depletion via dCTP deamination prompted us to ask whether there are additional phage resistance mechanisms that defend the cell by depleting deoxynucleotides. Depletion of the nucleotide pool was reported as an antiviral strategy in the human cell-autonomous innate immune system, where it is manifested by the SAMHD1 antiviral protein that restricts HIV infection in non-replicating cells (Goldstone et al., 2011). SAMHD1 was found to remove the triphosphate from dNTPs, breaking the nucleotides into phosphate-free deoxynucleosides and an inorganic triphosphate (PPPi) (Goldstone et al., 2011). The massive degradation of dNTPs by SAMHD1 depletes the nucleotide pool of the cells and inhibits replication of the viral genome (Ayinde et al., 2012).

Several dNTP triphosphohydrolases (dNTPases), and specifically dGTPases, have been previously described in bacteria (Kondo et al., 2007; Mega et al., 2009; Quirk and Bessman, 1991). Bacterial dGTPases cleave dGTP *in vitro*, and their structures show substantial homology to the active site architecture of SAMHD1 (Barnes et al., 2019; Singh et al., 2015). Although dGTPase homologs are abundant in bacteria, their physiological roles remain mostly unknown (Singh et al., 2015). We hypothesized that some bacterial dGTPases may play a role in defense against phages via dGTP depletion.

We identified multiple genes with predicted dGTPase domains that were localized within operons of type I restriction-modification systems, or near other known defense systems, suggesting they might have a role in phage defense (Doron et al., 2018) (Figure 3A-3B). These genes could not be directly aligned to the human SAMHD1, but a homology search via the HHpred tool (Zimmermann et al., 2018) showed significant similarity to the structure of SAMHD1. To check whether these genes confer phage resistance, we cloned six such dGTPases into *E. coli* MG1655. Following infection by a diverse set of phages, we found that four of the cloned genes substantially protected the *E. coli* host against multiple phages, with the most prominent defense observed for the dGTPase from *Shewanella putrefaciens* CN-32 (*Sp*-dGTPase) (Figure 3C; Figure S5; Table S5). A D101A point mutation in the histidine/aspartate (HD) motif in the predicted active site of *Sp*-dGTPase abolished defense, suggesting that the dGTPase functionality is essential for antiphage activity (Figure 3D).

**Figure 3.**
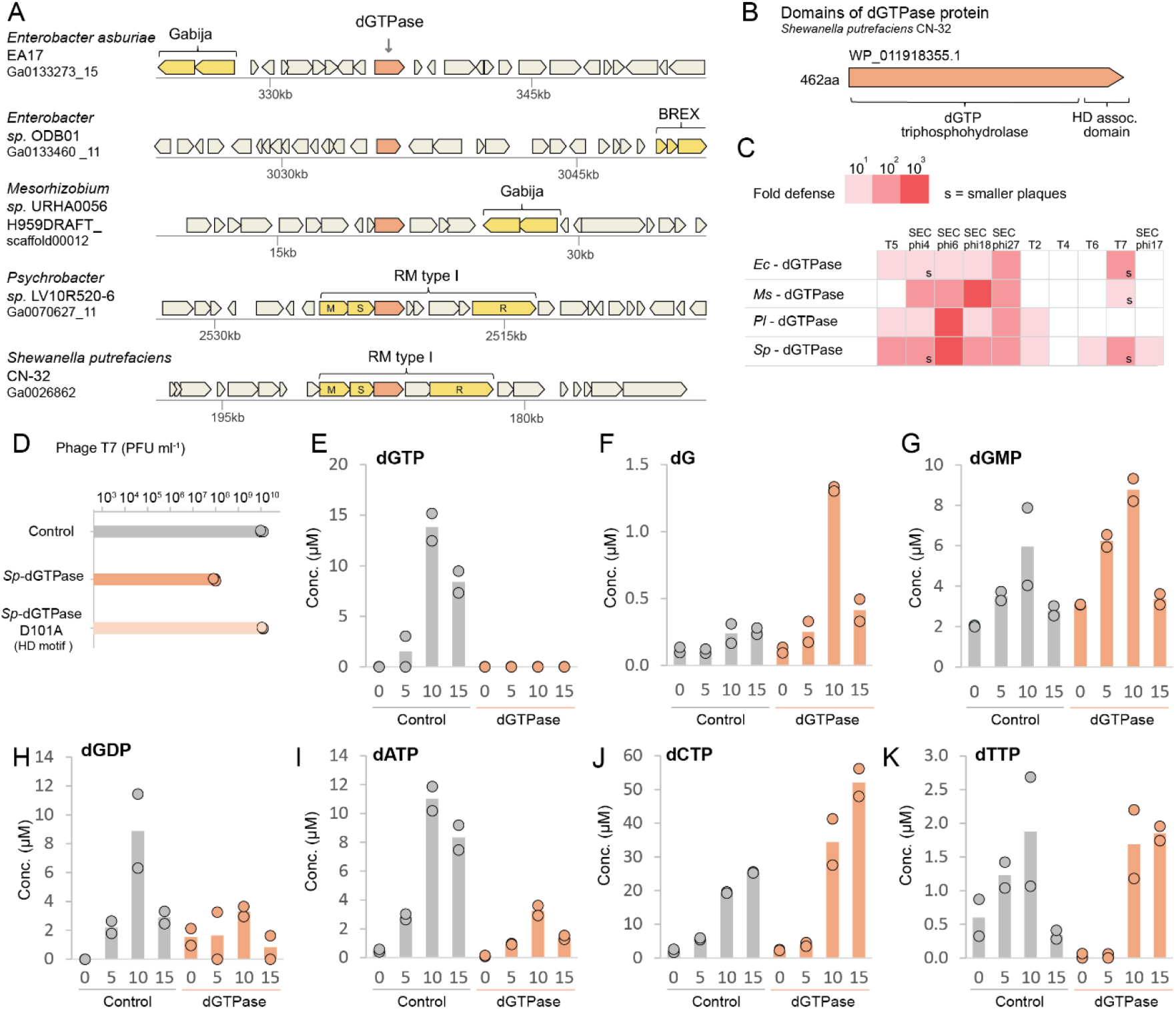
dGTPases defend against phages by depleting dGTP during phage infection. (A) Representative instances of dGTPase genes (in orange) and their genomic neighborhoods. Colors and annotations are as in Figure 1A. M, S and R designations within type I RM operon genes represent the methylase, specificity, and restriction subunits, respectively. (B) Domain organization of *Sp*-dGTPase. Protein accession in NCBI is indicated above the gene. (C). Multiple dGTPases defend against phages. Each dGTPase was cloned into a pBAD vector under the control of an arabinose-inducible promoter, and transformed into *E. coli* MG1655. Fold protection was measured using serial dilution plaque assays, comparing the efficiency of plating (EOP) of phages on the dGTPase-containing strain to the EOP on a control strain that lacks the gene. Data represent the average of three replicates (see expanded data in Figure S5). The designation ‘s’ stands for a marked reduction in plaque size. *Ec*, *E. coli* G177; *Ms*, *Mesorhizobium sp*. URHA0056; *Pl*, *Pseudoalteromonas luteoviolacea* DSM6061; *Sp*, *Shewanella putrefaciens* CN-32. (D) Plating efficiency of phage T7 on control cells, *Sp*-dGTPase-expressing cells, and a strain mutated in the predicted HD motif. Data represent plaque-forming units per ml (PFU/ml) in three replicates. (E-K) Concentrations of deoxynucleotides in cell lysates extracted from T7-infected cells, as measured by LC-MS with synthesized standards. X-axis represents minutes post infection, with zero representing non-infected cells. Cells were infected by phage T7 at an MOI of 2. Each panel shows data acquired for *Sp*-dGTPase-expressing cells or for control cells that contain an empty vector. Bar graphs represent the average of two biological replicates, with individual data points overlaid.

We next used LC-MS to analyze the nucleotide content in cells expressing *Sp*-dGTPase, which conferred strong defense against phage T7 (Figure 3C; Figure S5). In control cells that do not express the defensive gene, the concentration of dGTP was substantially elevated after 10 minutes from the onset of infection by phage T7, as expected for this phage (Lee and Richardson, 2011) (Figure 3E; Methods). However, in cells expressing the defensive gene, dGTP was undetectable throughout the infection time course (Figure 3E). In parallel to the depletion of dGTP, we observed substantial elevation in the concentrations of deoxyguanosine (dG), consistent with the hypothesis that the defensive protein removes the triphosphate from dGTP (Figure 3F). The monophosphorylated and diphosphorylated forms, dGMP and dGDP, were not similarly depleted, suggesting that the defensive dGTPase only degrades triphosphorylated guanosine forms (Figure 3G-H). These results reveal a bacterial antiviral enzyme which depletes dGTP in phage-infected cells and thus likely prevents phage replication.

The other triphosphorylated nucleotides (dCTP, dATP and dTTP) were not depleted in cells expressing *Sp*-dGTPase, suggesting that its primary substrate is dGTP (Figure 3I-3K). While we saw an elevation in dCTP concentrations in infected cells expressing the defensive dGTPase at the late stages of infection, the levels of dATP were somewhat reduced and those of dTTP remained unchanged, implying that perhaps the dGTPase targets dATP as a secondary substrate (Figure 3J-3K).

### Phages can evolve to escape nucleotide depletion

To gain more insight on the elements within phage infection that trigger nucleotide-depletion defense mechanisms, we attempted to isolate phage mutants that escape defense. We were able to obtain four mutants of phage T7 that could partially overcome the defense conferred by the dCTP deaminase gene, as well as seven mutants of phage T7 that overcame dGTPase defense, either partially or fully. We then sequenced the full genome of each of the mutants and compared the resulting sequences to the sequence of the wild-type phage (Supplementary Table S4).

In all four mutants that escaped dCTP deaminase defense, there were point mutations in gene 5.7 of phage T7, leading to amino-acid substitutions. These included H38R (mutants 1 and 4), C68Y (mutant 2), and R24W (mutant 3) (Figure 4A). These results suggest that mutations that alter the Gp5.7 protein of T7 enable the phage to overcome the defensive activity of the dCTP deaminase. Gp5.7 of phage T7 is responsible for shutting down σ^S^-dependent host RNA polymerase transcription, which would have otherwise interfered with phage propagation (Tabib-Salazar et al., 2018). Phages deleted in gene 5.7 were reported to be viable but propagate sub-optimally on *E. coli* cells (Tabib-Salazar et al., 2017); indeed, our mutant phages generate smaller plaques than the WT T7 (Figure 4B). It was previously shown that a point mutation in the arginine at position 24 of Gp5.7 abolishes its activity (Tabib-Salazar et al., 2017). In our mutant #3 this exact residue is mutated, suggesting that the function of Gp5.7 is likely impaired in this mutant (Figure 4A-4B).

**Figure 4.**
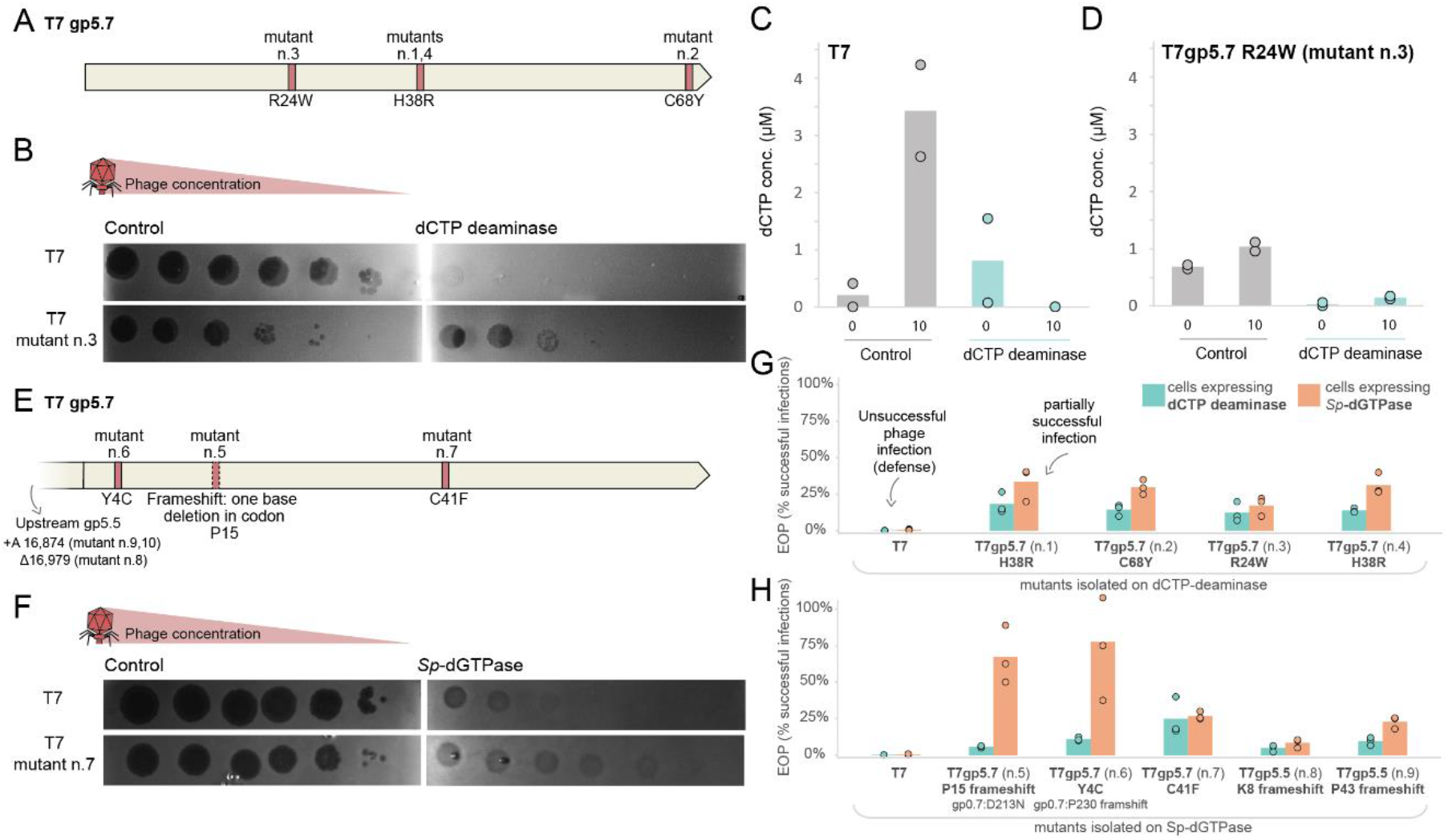
Phage mutants can overcome nucleotide-depletion defense. (A) Positions in the T7 gene 5.7 that are mutated in phages that escape defense by dCTP deaminase. (B) A representative phage mutant capable of escaping dCTP deaminase defense. Shown are ten-fold serial dilution plaque assays, comparing the plating efficiency of WT and mutant phages on bacteria that contain the dCTP deaminase and a control strain that lacks the system and contains an empty vector instead. Images are representative of three replicates. (C-D) Concentrations of dCTP in cell lysates extracted from cells infected by T7 (C) or T7 mutant #3 (D), as measured by LC-MS with synthesized standards. X-axis represents minutes post infection, with zero representing non-infected cells. Cells were infected by phage T7 at an MOI of 2. Each panel shows data acquired for dCTP deaminase-expressing cells or for control cells that contain an empty vector. Bar graphs represent average of two biological replicates, with individual data points overlaid. (E) Positions in the T7 gene 5.7 that are mutated in phages that escape defense by *Sp*-dGTPase. (F) A representative phage mutant capable of escaping *Sp*-dGTPase. (G) Efficiency of plating (EOP) of phage mutants that were isolated on dCTP deaminase cells. EOP is represented as the ratio between phage PFUs obtained on defense-containing cells and PFUs obtained on the control cells. Shown are EOP values when these mutants infect dCTP-deaminase-expressing cells (turquoise) or *Sp*-dGTPase-expressing cells (orange). (H) Same as panel G, for escape mutants isolated on *Sp*-dGTPase cells. The full list of mutations for each phage in this figure is detailed in Table S4.

We infected dCTP deaminase-expressing cells with the Gp5.7-mutated T7 phage, and measured dCTP concentrations. A reduction in dCTP was observed 10 minutes post infection, but this reduction caused only partial, yet not full, depletion of dCTP in the infected cells (Figure 4C-4D). These results suggest that the dCTP deaminase is only partially active against the mutated phage and that the remaining dCTP in the cell still allows phage DNA replication, explaining the ability of the phage to escape from defense.

We next examined the T7 mutants that escaped the *Sp*-dGTPase defense. Surprisingly, gene 5.7, the same gene that was mutated in the phages that overcame the dCTP deaminase, was also mutated in three of the seven T7 mutants that evaded *Sp*-dGTPase (Figure 4E-4F). In an additional three mutants, a frameshift-causing mutation was observed in gene 5.5, which is directly upstream to gene 5.7 and is thought to be translated together with gene 5.7, forming a 5.5-5.7 fusion protein (Dunn et al., 1983). Therefore, these mutations possibly render Gp5.7 non-translated (Figure 4E-4F). These mutations, which introduced mismatches, premature stop codons or frameshifts within the Gp5.7 protein, suggest that inactivation of the phage Gp5.7 protein allows phage T7 to escape dGTPase-based defense.

Some of the T7 mutants that escaped the *Sp*-dGTPase defense contained, in addition to the mutation in T7 gene 5.7, also mutations in other genes. Specifically, two escapees were also mutated in T7 gene 0.7 (Figure 4H, Supplementary Table S4). Gp0.7 encodes a phage protein kinase that phosphorylates the σ^70^-dependent host RNA polymerase, leading to increased transcription termination at sites found between the early and middle genes on the T7 genome (Severinova and Severinov, 2006). Notably, whereas T7 strains with a single mutation in gene 5.7 only partially escaped *Sp*-dGTPase defense, strains with an additional mutation in gene 0.7 escaped fully or nearly fully (Figure 4G-4H). These results suggest that mutations altering the capability of the phage to modify host RNA polymerase also enable the phage to overcome both the dCTP deaminase and *Sp*-dGTPase. A single T7 escapee was not mutated in either of these genes, but had a point mutation in the endonuclease I gene, encoding a protein responsible for Holliday junction resolution (Hadden et al., 2007) (Supplementary Table S4).

Phage escape mutants in gene 5.7, which were isolated on the dCTP deaminase-expressing strain, were also able to overcome defense by the dGTPase and vice versa, showing that the same mutations leads the T7 phage to resist defense by both enzymes (Figure 4G-4H). These results suggest that, although the two nucleotide-depletion mechanisms that we studied are enzymatically different from one another and do not share sequence or functional homology, both defensive genes likely sense the same pathway within the T7 infection cycle as a trigger that activates their dNTP depletion capacity (see Discussion).

### Numerous species encode antiviral nucleotide-depletion genes

We found homologs of the defensive dCTP deaminase in 952 genomes, representing 2.5% of the set of 38,167 genomes that we analyzed (Supplementary Table S1). The dCTP deaminases were found in genomes of a diverse set of species from the Proteobacteria phylum, as well as other phyla including Bacteroidetes, Actinobacteria, and Firmicutes (Figure 5A; Supplementary Table S1). In most cases, only a small fraction of the sequenced genomes of a given species harbored the dCTP deaminase. For example, the gene was found in 5%, 10%, and 12% of analyzed *E. coli*, *Legionella pneumophila*, and *Burkholderia pseudomallei* genomes, respectively (Figure 5B). Such a sparse pattern of gene presence/absence is typical of bacterial defense systems and is indicative of extensive horizontal gene transfer between genomes (Bernheim and Sorek, 2020). An exception for this pattern was observed in *Vibrio cholerae*, where we found the defensive dCTP deaminase in most of the analyzed genomes (200 of 291, 69%) (Figure 5B).

**Figure 5.**
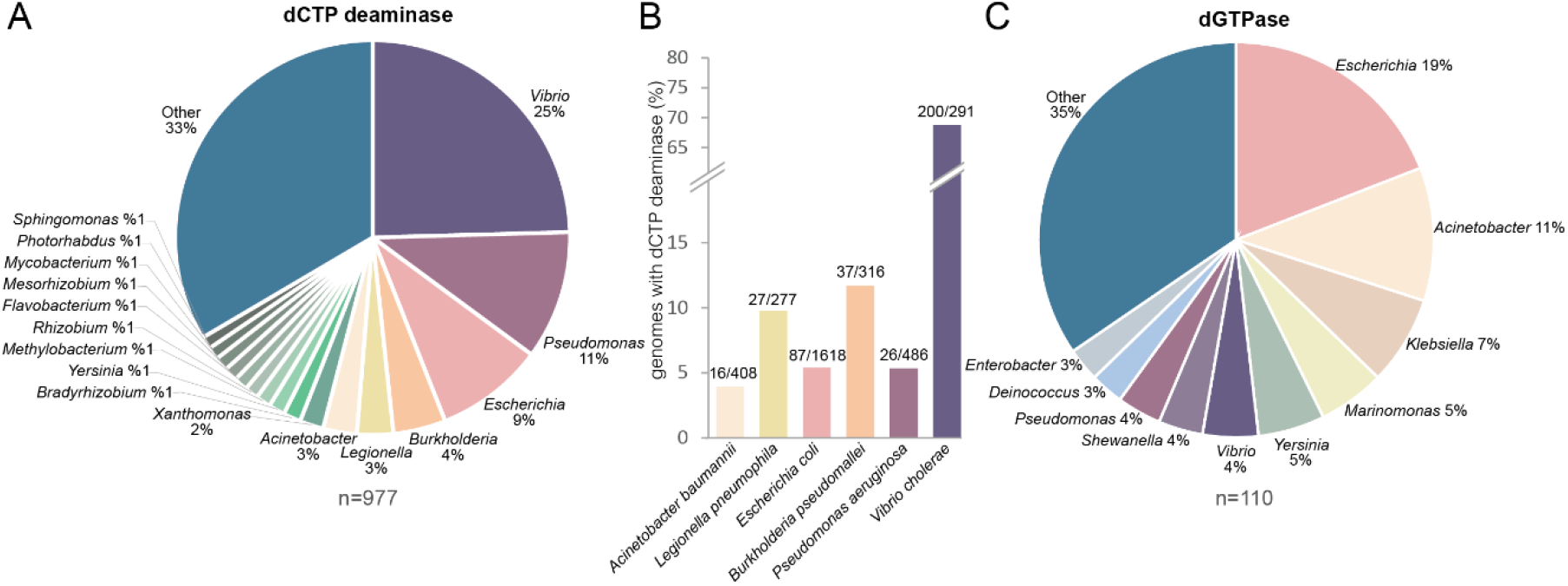
Distribution of homologs of nucleotide-depleting defense genes in bacterial genomes. (A) Distribution of 977 homologs of dCTP deaminase among bacterial genomes in the analyzed dataset. (B) Presence of dCTP deaminase homologs in selected species. The values above each bar represent the number of sequenced genomes that contain the dCTP deaminase homolog out of the total number of genomes of the species present in the analyzed database. (C) Distribution of 110 close homologs of *Sp*-dGTPases among bacterial genomes in the analyzed dataset.

Homologs of the defensive *Sp*-dGTPase were abundant in the set of bacterial genomes that we analyzed, appearing in >2,300 of the genomes (at least 25% sequence identity to *Sp*-dGTPase over an alignment overlap of >= 90%) (Supplementary Table S2). Among these, a set of 110 proteins showed high similarity (>=45% sequence identity) to *Sp*-dGTPase, and 53% of these (58/110) were localized next to known defense systems such as restriction enzymes and abortive infection genes (Supplementary Table S2). These results imply that the main role of these direct homologs is to defend against phages. The defensive homologs of *Sp*-dGTPase were found in a diverse set of Proteobacterial species, notably from the genera *Escherichia*, *Acinetobacter*, *Klebsiella*, and *Vibrio* (Figure 5C).

Homologs with sequence identity lower than 45% as compared to *Sp*-dGTPase did not show high propensity to co-localize with defense systems, with only 10% of these homologs found next to known defensive genes (Supplementary Table S2). These findings suggest that these more distant homologs may have non-defensive, possibly housekeeping, functions. Alternatively, it is possible that these homologs do function in phage defense, but also have housekeeping roles in bacterial physiology that prevent them from being carried on mobile genetic elements that transport defense systems between genomes.

## Discussion

In this study we describe a previously unknown mechanism of phage resistance, in which phage infection triggers the activity of defensive nucleotide-manipulating enzymes that deplete one of the deoxy nucleotides from the infected cell. As phage DNA replication necessitates a vast supply of all dNTPs, elimination of one of the dNTPs from the nucleotide pool would impair the phage ability to replicate its genome. It was previously shown that starvation for a single dNTP can generate abnormally branched DNA replication intermediates, and, if prolonged, can lead to bacterial cell death (Itsko and Schaaper, 2014). Therefore, depletion of a dNTP during phage infection may not only halt phage replication, but could eventually lead to abortive infection cell death, thus protecting the culture from phage propagation (Lopatina et al., 2020).

Enzymes with cytidine deaminase activity are known to be involved in multiple aspects of immunity (Harris and Dudley, 2015). The human APOBEC3G protects against viruses by deaminating cytosines in the viral genome, thus destroying its coding capacity (Okada and Iwatani, 2016; Stavrou and Ross, 2015). In addition, cytosine deamination performed by activation-induced cytidine deaminase (AID) enzymes introduces somatic hypermutations which are essential for antibody diversification and maturation (Kumar et al., 2014). We now report a third role for cytidine deaminases in immunity, where the deamination is performed on the mononucleotide building blocks in order to eliminate them during viral infection. Notably, the mechanisms of some vertebrate immune proteins that have predicted cytidine deaminase domains are still unknown (Harris and Dudley, 2015). It is tempting to hypothesize that some of these would function similarly to the bacterial defensive dCTP deaminase.

The model *E. coli* strain MG1655 encodes a dGTPase gene called *dgt*, which shows distant homology (24% identity) to the defensive *Sp*-dGTPase studied here (Wurgler and Richardson, 1990). The function of *dgt* was linked to maintenance of genome integrity (Gawel et al., 2008), although some evidence hints at a role in phage defense, as T7 was shown to encode a specific inhibitor of *dgt* (Myers et al., 1987). It is therefore possible that families of dGTPase enzymes other than the one that we studied here are also involved in phage defense. We note, however, that closely-related homologs of *dgt* are not preferentially localized within defense islands.

Multiple proteins in T7 modify the host RNA polymerase (RNAP) to allow optimal infection. These include Gp0.7, Gp2, and Gp5.7. Gp0.7 is a kinase that phosphorylates σ^70^-bound host RNAP and alters its functionality to suit phage needs (Tabib-Salazar et al., 2018). At later stages of infection, Gp2 shuts off the ability of σ^70^-RNAP to transcribe the phage genome (Tabib-Salazar et al., 2018). Finally, Gp5.7 binds and inhibits transcription by σ^S^-RNAP, which appears in the host when the cell activates the stringent response during phage infection (Tabib-Salazar et al., 2018). It is intriguing that mutations in two of these RNAP-modifying proteins allow the phage to escape both mechanisms of nucleotide depletion that are characterized here (notably, Gp2 is indispensable for T7 growth and hence cannot be mutated). This implies that both the dCTP deaminase and dGTPase somehow monitor the integrity of the host RNAP and become activated when it is tampered with.

Whereas we characterized two mechanisms for nucleotide depletion as a phage defense modality, it is likely that additional such mechanisms exist as part of the microbial antiviral arsenal. Enzymes that degrade all four dNTPs are known to exist in multiple bacteria including *Thermus thermophilus*, *Enterococcus faecalis*, and *Pseudomonas aeruginosa* (Kondo et al., 2008; Mega et al., 2009; Quirk and Bessman, 1991). These enzymes, which have *in vitro* activities similar to SAMHD1, may also play a role in anti-phage defense. Moreover, nucleotide-depleting antiviral proteins may have enzymatic reactions other than nucleotide deamination or triphosphohydrolysis. We envision that nucleotide methylases, ribosyltransferases, or other nucleotide modifying enzymes will be discovered in the future as having antiviral activities via manipulation of the nucleotide pool.

In the past few years, multiple components of the human innate immune system were found to have evolutionary roots in bacterial defense against phages. These include the cGAS-STING pathway (Morehouse et al., 2020), the viperin antiviral protein (Bernheim et al., 2021), the argonaute protein of the RNAi machinery (Kuzmenko et al., 2020), and TIR domains (Ofir et al., 2021). The defensive dGTPases characterized here show distant structural homology to the SAMHD1 active site, but this homology is too limited to determine whether the SAMHD1 is evolutionarily derived from a bacterial dNTPase. Regardless of the evolutionary trajectory, we find it remarkable that depletion of the nucleotide pool is a successful antiviral defense strategy shared by eukaryotes and prokaryotes alike.

## Acknowledgements

We would like to thank Pascale Cossart for pointing us to the SAMHD1 antiviral mechanism, and Aude Bernheim and the Sorek laboratory members for comments on earlier versions of this manuscript and fruitful discussion. R.S. was supported, in part, by the European Research Council (grant ERC-CoG 681203), the Ernest and Bonnie Beutler Research Program of Excellence in Genomic Medicine, the Minerva Foundation with funding from the Federal German Ministry for Education and Research, the Knell Family Center for Microbiology, the Yotam project and the Weizmann Institute Sustainability And Energy Research (SAERI) initiative, and the Dr. Barry Sherman Institute for Medicinal Chemistry. A.M. was supported by a fellowship from the Ariane de Rothschild Women Doctoral Program and, in part, by the Israeli Council for Higher Education via the Weizmann Data Science Research Center, and by a research grant from Madame Olga Klein-Astrachan.

## Methods

### Detection of nucleotide-depleting systems in defense islands

Protein sequences of all genes in 38,167 bacterial and archaeal genomes were downloaded from the Integrated Microbial Genomes (IMG) database (Chen et al., 2019) in October 2017. These proteins were filtered for redundancy using the ‘clusthash’ option of MMseqs2 (release 2-1c7a89) (Steinegger and Söding, 2017) using the ‘-min-seq-id 0.9’ parameter and then clustered using the ‘cluster’ option, with default parameters. Each cluster with >10 genes was annotated with the most common pfam, COG and product annotations in the cluster. Defense scores were calculated as previously described (Doron et al., 2018), recording the fraction of genes in each cluster that have known defense genes in their genomic environment spanning 10 genes upstream and downstream the inspected gene. In addition, each cluster was processed using Clustal-Omega (version 1.2.4) (Sievers et al., 2011) to produce a multiple sequence alignment. The alignment of each cluster was searched using the ‘hhsearch’ option of hhsuite (version 3.0.3) (Soding, 2005) against the PDB70 and pfamA_v32 databases, using the ‘−p 10 −loc −z 1 −b 1 −ssm 2 −sc 1 −seq 1 – dbstrlen 10000 −maxres 32000 −M 60 −cpu 1’ parameters. Clusters that had HHsearch hits (Zimmermann et al., 2018) both to an N-terminal kinase and a C-terminal dCTP deaminase, sized >350aa, and which had a defense score >0.33 were included in the family of the dCTP deaminase genes studied here. Clusters sized <=10 genes, whose representative sequence (Steinegger and Söding, 2017) aligned to the representative sequences of selected dCTP deaminase clusters with an e-value <0.01, and whose size was >350aa, were also included in the family, adding 30 genes to the list in Supplementary Table S1. To generate the list of dGTPase homologs in Supplementary Table S2 we aligned the studied *Sp*-dGTPase protein to all proteins in the database using the ‘search’ option of MMseqs2 (release 12-113e3) using the ‘−s 7 −threads 20 −max-seqs 100000’ parameters, and retained all hits sized >400aa that had >=25% sequence identity to *Sp*-dGTPase, with alignment overlap of at least 90% on both query and subject sequences.

### Bacterial strains and phages

*E. coli* strain MG1655 (ATCC 47076) was grown in MMB (LB + 0.1 mM MnCl2 + 5 mM MgCl2, with or without 0.5% agar) at 37 °C or room temperature (RT). Whenever applicable, media were supplemented with ampicillin (100 μg ml−1), to ensure the maintenance of plasmids. Infection was performed in MMB media at 37°C or RT as detailed in each section. Phages used in this study are listed in Supplementary Table S3.

### Plasmid and strain construction

dCTP deaminase and dGTPase genes used in this study were synthesized by Genscript Corp. and cloned into the pSG1 plasmid (Doron et al., 2018) with their native promoters, or into the pBad plasmid (Thermofisher, cat. #43001), respectively, as previously described (Bernheim et al., 2021; Doron et al., 2018). Mutants of the dCTP deaminase gene of *E. coli* AW1.7 were also synthesized and cloned by Genscript. All synthesized sequences are presented in Table S5. Mutant of the dGTPase gene of *Shewanella putrefaciens* CN-32 was constructed using Q5 Site directed Mutagenesis kit (NEB, cat. #E0554S) using primers presented in Supplementary Table S5.

### Plaque assays

Phages were propagated by picking a single phage plaque into a liquid culture of *E. coli* MG1655 grown at 37°C to OD_600_ of 0.3 in MMB medium until culture collapse. The culture was then centrifuged for 10 minutes at 4000 rpm and the supernatant was filtered through a 0.2 μm filter to get rid of remaining bacteria and bacterial debris. Lysate titer was determined using the small drop plaque assay method as described before (Mazzocco et al., 2009).

Plaque assays were performed as previously descried (Mazzocco et al., 2009). Bacteria (*E. coli* MG1655 with pSG1-dCTP deaminase or pBad-dGTPase) and negative control (*E. coli* MG1655 with empty pSG1 or pBad-GFP) were grown overnight at 37°C. Then 300 μl of the bacterial culture was mixed with 30 ml melted MMB agar (LB + 0.1 mM MnCl2 + 5 mM MgCl2 + 0.5% agar, with or without 0.2% arabinose, as described in Table S5) and left to dry for 1 hour at room temperature. 10-fold serial dilutions in MMB were performed for each of the tested phages and 10 μl drops were put on the bacterial layer. Plates were incubated overnight at RT. Plaque forming units (PFUs) were determined by counting the derived plaques after overnight incubation.

### DNA and RNA editing assays

Overnight cultures of bacteria (*E. coli* MG1655 harboring pSG1-dCTP deaminase plasmid) or negative control (*E. coli* MG1655 with the pSG1 plasmid) were diluted 1:100 in 60 ml of MMB medium and incubated at 37 °C while shaking at 200 rpm until early log phase (OD_600_ of 0.3). 10 ml samples of each bacterial culture were taken and centrifuged at 4000 rpm for 5 min at 4 °C. The pellets were flash frozen using dry ice and ethanol. The remaining cultures were infected by phage T7 at a final MOI of 2. 10 ml samples were taken throughout infection at 5, 10 and 15 minutes post infection, and centrifuged and flash frozen as described above.

DNA of all samples was extracted using the QIAGEN DNeasy blood and tissue kit (cat #69504), using the gram negative bacteria protocol. Libraries were prepared for Illumina sequencing using a modified Nextera protocol as previously described (Baym et al., 2015). Following sequencing on an Illumina NextSeq500 the sequence reads were aligned to bacterial and phage reference genomes (GenBank accession numbers: NC_000913.3, NC_001604.1, respectively) by using NovoAlign (Novocraft) v3.02.02 with default parameters, and mutations were identified and quantified by counting each mismatch across the genome. Frequency of mismatches was compared between control and dCTP deaminase samples throughout the infection time course.

RNA extraction was performed as described previously (Dar et al., 2016). Briefly, frozen pellets were re-suspended in 1 ml of RNA protect solution (FastPrep) and lysed by Fastprep homogenizer (MP Biomedicals). RNA was extracted using the FastRNA PRO blue kit (MP Biomedicals, 116025050) according to the manufacturer’s instructions. DNase treatment was performed using the Turbo DNA free kit (Life Technologies, AM2238). RNA was subsequently fragmented using fragmentation buffer (Ambion-Invitrogen, cat. #10136824) at 72°C for 1 min and 45 sec. The reactions were cleaned by adding ×2.5 SPRI beads (Agencourt AMPure XP, Beckman-Coulter, A63881). The beads were washed twice with 80% ethanol and air dried for 5 minutes. The RNA was eluted using H_2_O. Ribosomal RNA was depleted by using the Ribo-Zero rRNA Removal Kit (epicentre, MRZB12424). Strand-specific RNA-seq was performed using the NEBNext Ultra Directional RNA Library Prep Kit (NEB, E7420) with the following adjustments: all cleanup stages were performed using ×1.8 SPRI beads, and only one cleanup step was performed after the end repair step. Following sequencing on an Illumina NextSeq500, sequenced reads were demultiplexed and adapters were trimmed using ‘fastx clipper’ software with default parameters. Reads were mapped to the bacterial and phage genomes by using NovoAlign (Novocraft) v3.02.02 with default parameters, discarding reads that were non-uniquely mapped as previously described (Dar et al., 2016). Mutations from reference genomes were identified and quantified by counting each mismatch across the transcriptome. Frequency of mismatches was compared between control and dCTP deaminase samples throughout the infection time course.

### Cell lysate preparation

Overnight cultures of *E. coli* harboring the defensive gene and negative controls were diluted 1:100 in 250 ml MMB medium (with or without 0.2% arabinose, as described in Table S5) and grown at 37 °C (250 rpm) until reaching OD_600_ of 0.3. The cultures were infected by T7 at a final MOI of 2. Following the addition of phage, at 5, 10 and 15 minutes post infection (plus an uninfected control sample), 50 ml samples were taken and centrifuged for 5 minutes at 4000 rpm. Pellets were flash frozen using dry ice and ethanol. The pellets were re-suspended in 600 μl of 100 mM phosphate buffer at pH=8 and supplemented with 4 mg/ml lysozyme. The samples were then transferred to a FastPrep Lysing Matrix B 2 ml tube (MP Biomedicals cat. #116911100) and lysed using a FastPrep bead beater for 40 seconds at 6 m/s (two cycles). Tubes were then centrifuged at 4°C for 15 minutes at 15,000g. Supernatant was transferred to Amicon Ultra-0.5 Centrifugal Filter Unit 3 kDa (Merck Millipore cat. #UFC500396) and centrifuged for 45 minutes at 4°C at 12,000g. Filtrate was taken and used for LC-MS analysis.

### Quantification of nucleotides by HPLC-MS

Cell lysates were prepared as described above and sent for analysis at MS-Omics Ltd. Quantification of nucleotides and nucleosides was carried out using a Vanquish™ Horizon UHPLC System coupled to Q Exactive™ HF Hybrid Quadrupole-Orbitrap™ Mass Spectrometer (Thermo Fisher Scientific, US). The UHPLC was performed using an Infinity Lab PoroShell 120 HILIC-Z PEEK lined column with the dimension of 2.1 × 150mm and particle size of 2.7μm (Agilent Technologies). The composition of mobile phase A was 10 mM ammonium acetate at pH 9 in 90% Acetonitrile LC-MS grade (VWR Chemicals, Leuven) and 10% ultra-pure water from Direct-Q^®^ 3 UV Water Purification System with LC-Pak^®^ Polisher (Merck KGaA, Darmstadt) and mobile phase B was 10 mM ammonium acetate at pH 9 in ultra-pure water with 12 μM medronic acid (InfinityLab Deactivator additive, Agilent Technologies). The flow rate was kept at 250 μl/ml consisting of a 2 minutes hold at 10% B, increased to 40% B at 14 minutes, held until 15 minutes, decreased to 10% B at 16 minutes and held for 8 minutes. The column temperature was set at 30 °C and an injection volume of 5 μl. An electrospray ionization interface was used as ionization source. Analysis was performed in positive ionization mode from m/z 200 to 900 at a mass resolution of 120000.

Sample analysis was carried out by MS-Omics as follows. Samples and standards were diluted 1:3 in 10% ultra-pure water and 90% acetonitrile containing 10 mM ammonium acetate at pH 9, then filtered through a Costar^®^ Spin-X ^®^ centrifuge tube filter 0.22 μm nylon membrane. The majority of the standard used for quantification were purchased from Sigma Aldrich (Steinheim, Germany): adenosine, adenosine 5′-monophosphate sodium salt, adenosine 5′-diphosphate sodium salt, adenosine 5′-triphosphate disodium salt hydrate, cytidine, cytidine 5′-monophosphate disodium salt, cytidine 5′-diphosphate sodium salt hydrate, cytidine 5′-triphosphate disodium salt, guanosine, guanosine 5′-monophosphate disodium salt hydrate, guanosine 5′-diphosphate sodium salt, guanosine 5′-triphosphate sodium salt hydrate, uridine, uridine 5′-monophosphate disodium salt, uridine 5′-(trihydrogen diphosphate) sodium salt, uridine 5′-triphosphate trisodium salt hydrate, 2′-deoxyadenosine monohydrate, 2’-deoxyadenosine 5’-diphosphate sodium salt, 2′-deoxycytidine, 2’-deoxycytidine 5’-monophosphate, 2’-deoxycytidine 5’-diphosphate, 2’-deoxyguanosine, 2’-deoxyguanosine 5’-diphosphate, thymidine, thymidine 5’-monophosphate disodium salt hydrate, thymidine 5’-diphosphate disodium salt hydrate, 2′-deoxyuridine, 2′-deoxyuridine 5′-triphosphate sodium salt and a deoxynucleotides mix (10mM, PCR reagent) containing 2′-deoxyadenosine 5′-triphosphate, 2′-deoxycytidine 5′-triphosphate, 2′-deoxyguanosine 5′-triphosphate and thymidine 5′-triphosphate. Other standards were purchased through MetaSci (Toronto, Cananda): 2’-deoxyuridine, 2’-deoxyadenosine-5’-monophosphate and 2’-deoxyguanosine 5’-monophosphate sodium salt hydrate. Peak areas were extracted using Trace Finder™ Version 4.1 (Thermo Fisher Scientific, US) and quantified using a calibration ranging from 0.005 to 40 μM. Deoxyuridine monophosphate and diphosphate were quantified using the linear regression of uridine monophosphate and diphosphate, assuming a similar response factor.

### Isolation of mutant phages

To isolate mutant phages that escape dCTP deaminase and dGTPase defense, phages were plated on bacteria expressing each defense system using the double-layer plaque assay (Kropinski et al., 2009). Bacterial cells expressing each defense system were grown in MMB (supplemented with 0.2% arabinose for the dGTPase) to an OD_600_ of 0.3. Then, 100 μl of phage lysate was mixed with 100 μl bacterial cells expressing each defense gene, and left at room temperature for 10 minutes. 5 ml of pre-melted 0.5% MMB (supplemented with 0.2% arabinose for the dGTPase) was added and the mixture was poured onto a bottom layer of 1.1% MMB. For the dGTPase, the double layer plates were incubated overnight at 37°C, and single plaques were picked into 90 ml phage buffer. For the dCTP deaminase, double layer plates were incubated overnight at room temperature and the entire top layer was scraped into 2 ml of phage buffer to enrich for phages that escape the defense genes. Phages were left for 1 h at room temperature during which the phages were mixed several times by vortex to release them from the agar into the phage buffer. The phages were centrifuged at 3200 g for 10 min to get rid of agar and bacterial cells, and the supernatant was transferred to a new tube. For both defense genes, in order to test the phages for the ability to escape from the defense system, the small drop plaque assay was used (Mazzocco et al., 2009). 300 ml bacteria harboring the defensive gene or negative control (*E. coli* MG1655 with plasmid pSG1 or pBad-GFP lacking the system) were mixed with 30 mL melted MMB 0.5% agar and left to dry for 1 hour at room temperature. 10-fold serial dilutions in phage buffer were performed for the ancestor phages (WT phage used for the original double layer plaque assay) and the phages formed on the strain expressing the defense genes. 10 μl drops were put on the bacterial layer. The plates were incubated overnight at room temperature or 37°C for dCTP deaminase or dGTPase, respectively.

### Amplification of mutant phages

Isolated phages for which there was decreased defense compared to the ancestor phage were further propagated by picking a single plaque formed on the defense gene in the small drop plaque assay into a liquid culture of *E. coli* harboring the defensive gene, which was grown at 37°C in MMB with shaking at 200 rpm to an OD_600_ of 0.3. The phages were incubated with the bacteria at 37°C with shaking at 200 rpm for 3 hr, and then an additional 9 mL of bacterial culture grown to OD_600_ of 0.3 in MMB was added, and incubated for additional 3 h in the same conditions. For the dGTPase system the growth media was supplemented with 0.2% arabinose throughout the entire amplification process. The lysates were then centrifuged at 4000 rpm for 10 min and the supernatant was filtered through a 0.2 μm filter to get rid of remaining bacteria. Phage titer was then checked using the small drop plaque assay on the negative control strain as described above.

### Sequencing and genome analysis of phage mutants

High titer phage lysates (>10^7^ PFU/ml) of the ancestor phage and isolated phage mutants were used for DNA extraction. 0.5 ml of phage lysate was treated with DNase-I (Merck cat #11284932001) added to a final concentration of 20 ug/ml and incubated at 37°C for 1 hour to remove bacterial DNA. DNA was extracted using the QIAGEN DNeasy blood and tissue kit (cat. #69504) starting from a proteinase-K treatment to degrade the phage capsids. Libraries were prepared for Illumina sequencing using a modified Nextera protocol as described above. Following sequencing on Illumina NextSeq500, reads were aligned to the phage reference genome (GenBank accession number: NC_001604.1) and mutations compared to the reference genome were identified using Breseq (versions 0.29.0 or 0.34.1 for mutant phages that escape dCTP deaminase and dGTPase, respectively) with default parameters (Deatherage and Barrick, 2014). Only mutations that occurred in the isolated mutants, but not in the ancestor phage, were considered. Silent mutations within protein coding regions were disregarded as well.

## Supplementary Figures

**Figure S1.**
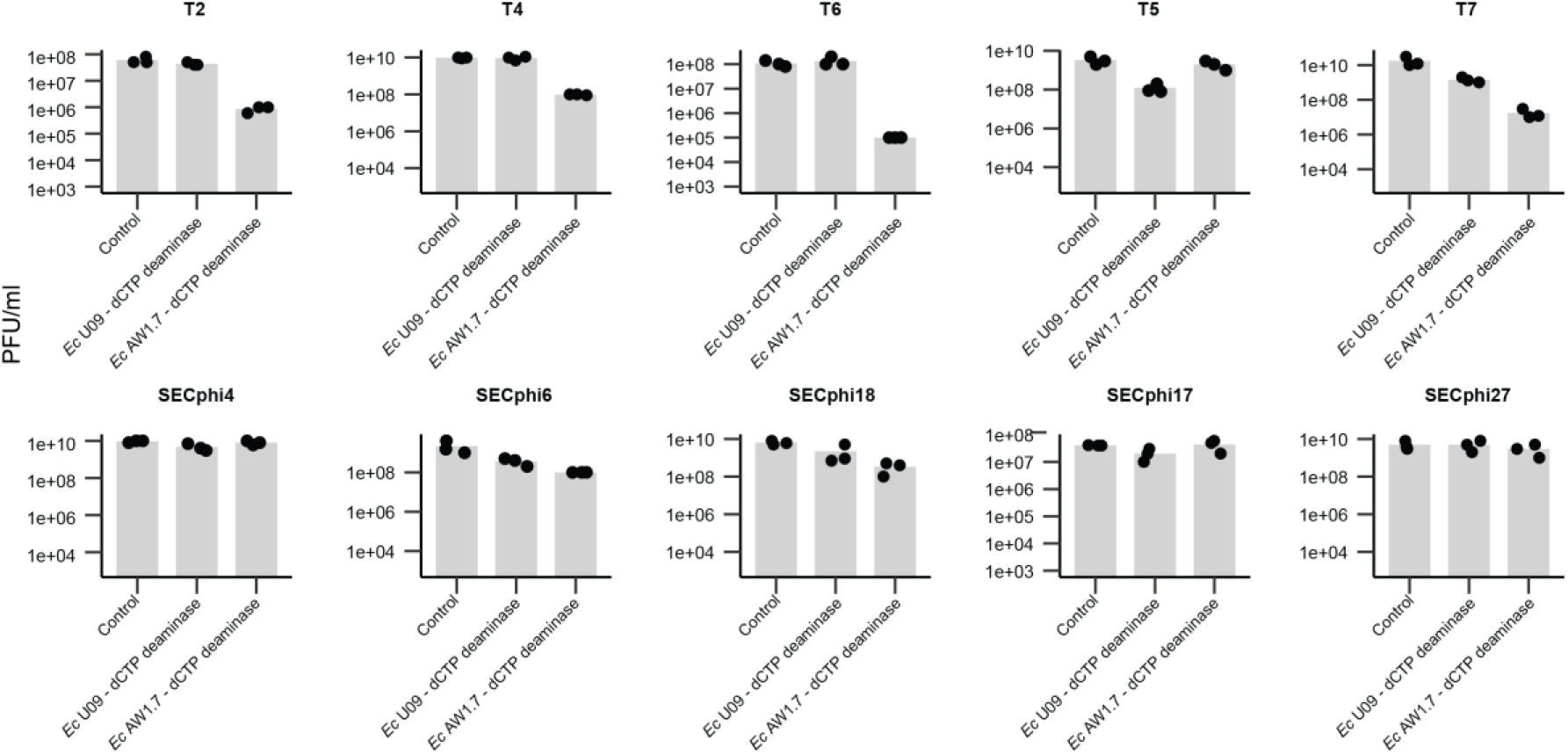
dCTP deaminases protect against phage infection. Bacteria expressing dCTP deaminases from *E. coli* U09 or *E. coli* AW1.7, as well as a negative control that contains an empty vector, were grown on agar plates in room temperature. Tenfold serial dilutions of the phage lysate were dropped on the plates. Data represent plaque-forming units per milliliter for ten phages tested in this study. Each bar graph represents average of three replicates, with individual data points overlaid.

**Figure S2.**
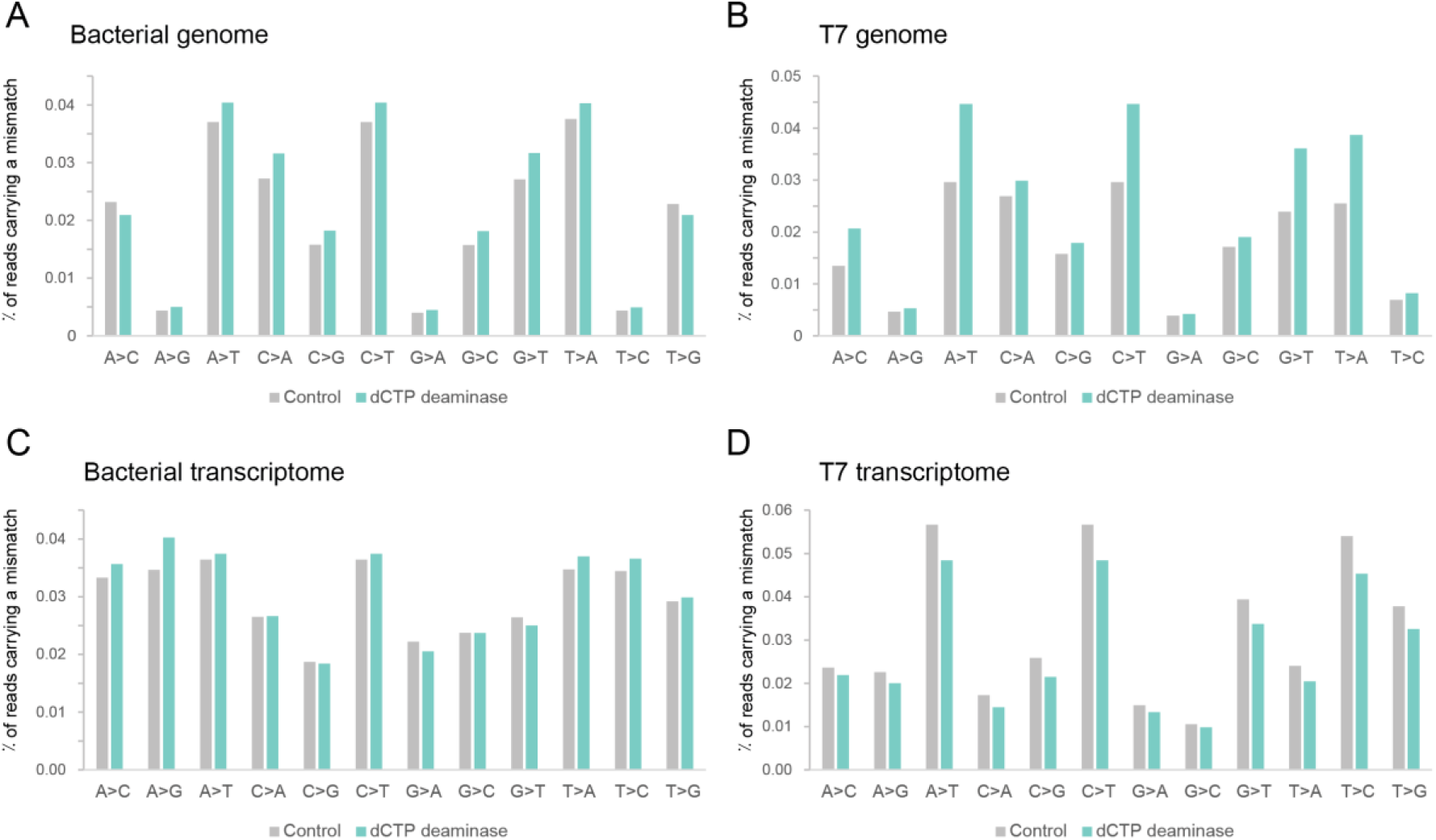
No evidence for editing of genome and transcriptome by the dCTP deaminase. Cells expressing the deaminase from *E. coli* AW1.7 were infected by phage T7 at a multiplicity of infection (MOI) of 2 at 37°C. Total DNA and total RNA were extracted after 15 minutes from the onset of infection, and were subjected to DNA-seq and RNA-seq, respectively. Panels A and B show the abundance of DNA reads with specific mismatches for reads aligned to the bacterial genome (A) or the phage genome (B). Panels C and D show the abundance of RNA-seq reads with specific mismatches for reads aligned to the bacterial genome (C) or the phage genome (D).

**Figure S3.**
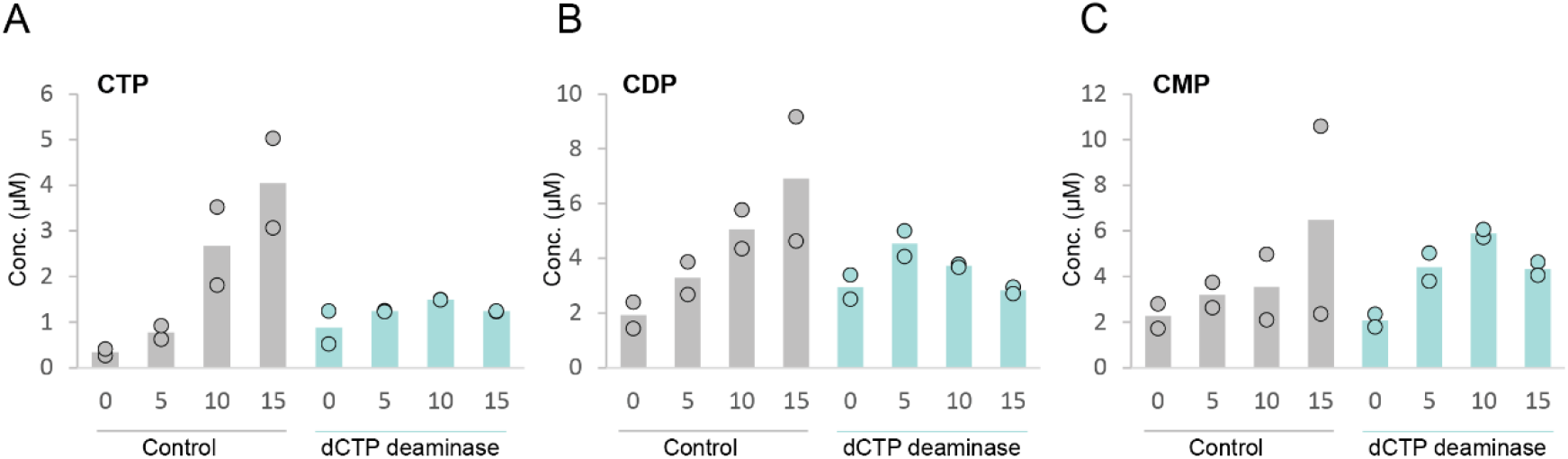
Cytidine ribonucleotides during phage infection. Concentrations of cytidine ribonucleotides in cell lysates extracted from T7-infected cells, as measured by LC-MS with synthesized standards. X axis represents minutes post infection, with zero representing non-infected cells. Cells were infected by phage T7 at an MOI of 2. Each panel shows data acquired for dCTP deaminase-expressing cells or for control cells that contain an empty vector. Bar graphs represent average of two biological replicates, with individual data points overlaid.

**Figure S4.**
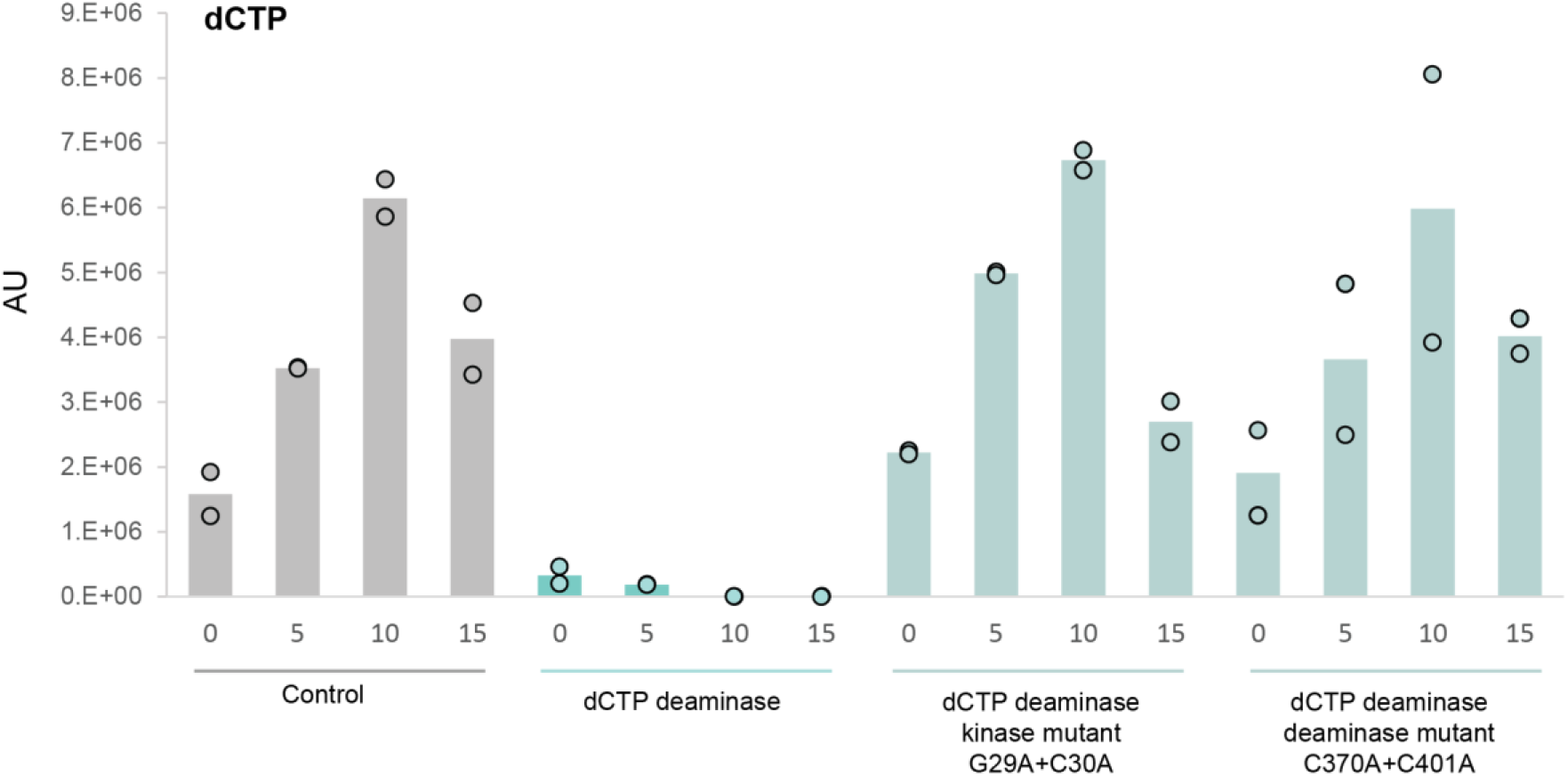
Mutated dCTP deaminase does not elicit dCTP depletion. Relative abundance of dCTP in cell lysates extracted from T7-infected cells, as measured by LC-MS. X axis represents minutes post infection, with zero representing non-infected cells. Y axis represents the area under the peak for dCTP, in arbitrary units (AU). Cells were infected by T7 at an MOI of 2. Presented are data acquired for cells expressing the dCTP deaminase from *E. coli* AW1.7, control cells that contain an empty vector, or cell expressing mutated forms of the dCTP deaminase. Bar graphs represent average of two biological replicates, with individual data points overlaid.

**Figure S5.**
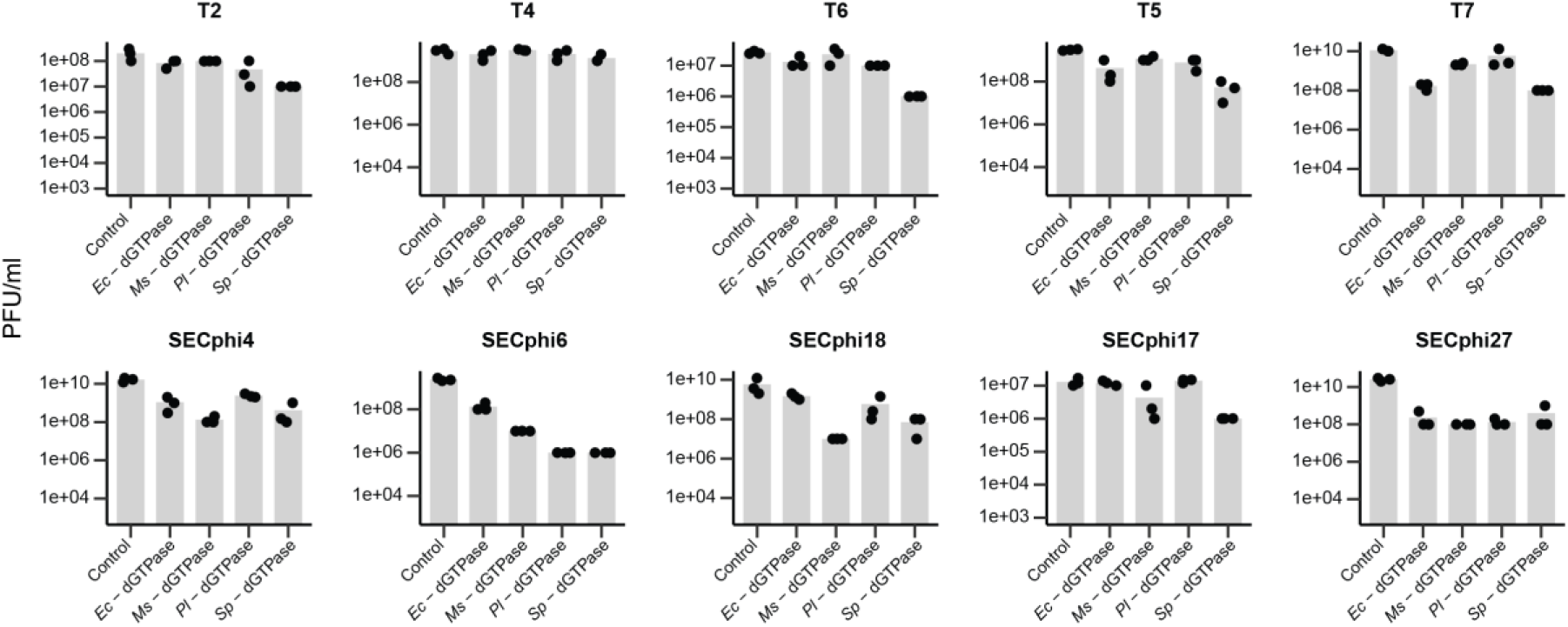
dGTPases protect against phage infection. *E. coli* MG1655 cells expressing dGTPases cloned from four different species (*Ec*, *E. coli* G177; *Ms*, *Mesorhizobium sp*. URHA0056; *Pl*, *Pseudoalteromonas luteoviolacea* DSM6061; *Sp*, *Shewanella putrefaciens* CN-32), as well as a negative control, were grown on agar plates in room temperature in the presence of 0.2% arabinose. Tenfold serial dilutions of the phage lysate were dropped on the plates. Data represent plaque-forming units per milliliter for ten phages tested in this study. Each bar graph represents average of three replicates, with individual data points overlaid.

## Supplementary Tables

Table S1. Homologs of the dCTP deaminases in this study (attached as an Excel spreadsheet).

Table S2. Homologs of the dGTPases in this study (attached as an Excel spreadsheet).

Table S3. Phages used in this study.

Table S4. Mutations observed in phages that escape nucleotide-depletion defense.

Table S5. Defense genes synthetized and cloned in this study (attached as an Excel spreadsheet).

Table S6. Primers used in this study.

**Table S3.** Phages used in this study.

| *Phage* | *Source* | *Identifier* | *Accession* |
| --- | --- | --- | --- |
| *Phage SECPhi17* | Doron et al., 2018 | N/A | LT960607.1 |
| *Phage SECPhi18* | Doron et al., 2018 | N/A | LT960609.1 |
| *Phage SECphi27* | Doron et al., 2018 | N/A | LT961732.1 |
| *Phage SECphi6* | Milman et al., 2020 | N/A | CADCZA010000001.1 |
| *Phage SECphi4* | Milman et al., 2020 | N/A | MT331608 |
| *Phage T2* | German Collection of Microorganisms and Cell Cultures GmbH (DSMZ) | DSM 16352 | LC348380.1 |
| *Phage T4* | U. Qimron | N/A | AF158101.6 |
| *Phage T5* | U. Qimron | N/A | AY543070.1 |
| *Phage T6* | German Collection of Microorganisms and Cell Cultures GmbH (DSMZ) | DSM 4622 | MH550421.1 |
| *Phage T7* | U. Qimron | N/A | NC_001604.1 |

**Table S4.**
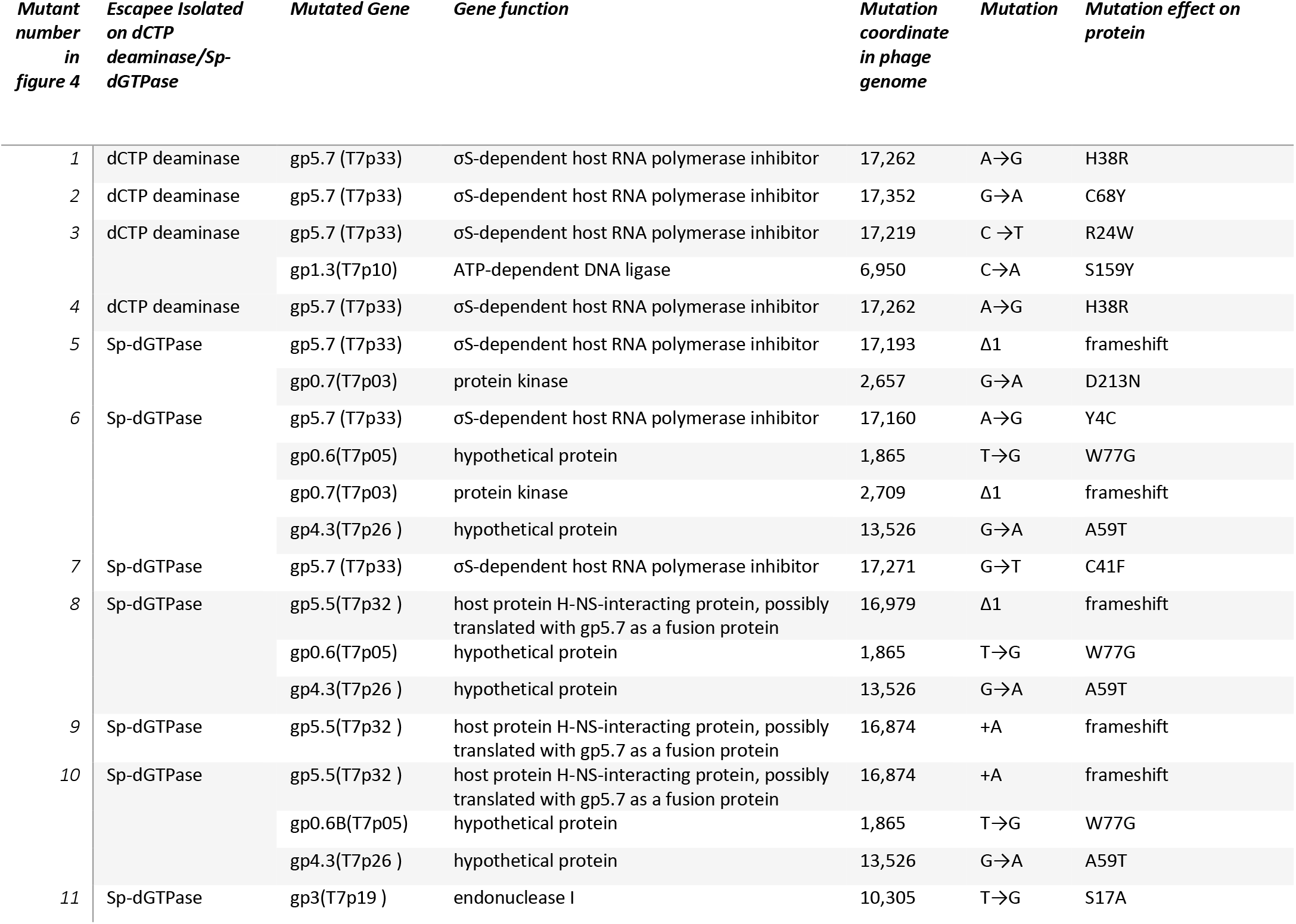
Mutations observed in phages that escape nucleotide-depletion defense.

| *Mutant number in figure 4* | *Escapee Isolated on dCTP deaminase/Sp-dGTPase* | *Mutated Gene* | *Gene function* | *Mutation coordinate in phage genome* | *Mutation* | *Mutation effect on protein* |
| --- | --- | --- | --- | --- | --- | --- |
| 1 | dCTP deaminase | gp5.7 (T7p33) | σS-dependent host RNA polymerase inhibitor | 17,262 | A→G | H38R |
| 2 | dCTP deaminase | gp5.7 (T7p33) | σS-dependent host RNA polymerase inhibitor | 17,352 | G→A | C68Y |
| 3 | dCTP deaminase | gp5.7 (T7p33) | σS-dependent host RNA polymerase inhibitor | 17,219 | C →T | R24W |
|  |  | gp1.3(T7p10) | ATP-dependent DNA ligase | 6,950 | C→A | S159Y |
| 4 | dCTP deaminase | gp5.7 (T7p33) | σS-dependent host RNA polymerase inhibitor | 17,262 | A→G | H38R |
| 5 | Sp-dGTPase | gp5.7 (T7p33) | σS-dependent host RNA polymerase inhibitor | 17,193 | Δ1 | frameshift |
|  |  | gp0.7(T7p03) | protein kinase | 2,657 | G→A | D213N |
| 6 | Sp-dGTPase | gp5.7 (T7p33) | σS-dependent host RNA polymerase inhibitor | 17,160 | A→G | Y4C |
|  |  | gp0.6(T7p05) | hypothetical protein | 1,865 | T→G | W77G |
| | | gp0.7(T7p03) | protein kinase | 2,709 | Δ1 | frameshift |
|  |  | gp4.3(T7p26 ) | hypothetical protein | 13,526 | G→A | A59T |
| 7 | Sp-dGTPase | gp5.7 (T7p33) | σS-dependent host RNA polymerase inhibitor | 17,271 | G→T | C41F |
| 8 | Sp-dGTPase | gp5.5(T7p32 ) | host protein H-NS-interacting protein, possibly translated with gp5.7 as a fusion protein | 16,979 | Δ1 | frameshift |
|  |  | gp0.6(T7p05) | hypothetical protein | 1,865 | T→G | W77G |
|  |  | gp4.3(T7p26 ) | hypothetical protein | 13,526 | G→A | A59T |
| 9 | Sp-dGTPase | gp5.5(T7p32 ) | host protein H-NS-interacting protein, possibly translated with gp5.7 as a fusion protein | 16,874 | +A | frameshift |
| 10 | Sp-dGTPase | gp5.5(T7p32 ) | host protein H-NS-interacting protein, possibly translated with gp5.7 as a fusion protein | 16,874 | +A | frameshift |
|  |  | gp0.6B(T7p05) | hypothetical protein | 1,865 | T→G | W77G |
|  |  | gp4.3(T7p26 ) | hypothetical protein | 13,526 | G→A | A59T |
| 11 | Sp-dGTPase | gp3(T7p19 ) | endonuclease I | 10,305 | T→G | S17A |

**Table S6.**
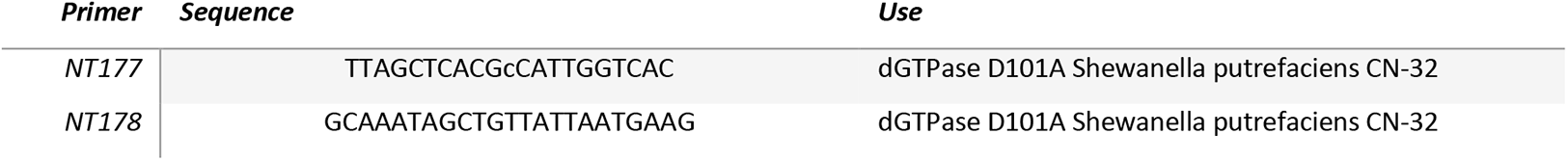
Primers used in this study.

